# Are cell length and volume interchangeable in cell cycle analysis?

**DOI:** 10.1101/2024.07.09.602793

**Authors:** Prathitha Kar, Ariel Amir

## Abstract

Cell length has been used as a proxy for cell size in cell cycle modeling studies. A previous study, however, brought into question the validity of this assumption, noting that correlations between cell lengths can be different from those involving cell volume if cell width fluctuations are taken into account. If cell volume is regulated, data analysis involving cell lengths will lead to an incorrect inference of the cell size control mechanism. We used conditional correlation of length variables conditioned upon radius variables to elucidate the underlying volume control mechanism. Using the conditional correlation on previous mother machine datasets measuring lengths at birth and division, and the cell radius for multiple cells, we find that the cell volume control strategy is consistent with an adder model. Further, using the conditional correlation, we conclude that measurement noise constitutes a significant portion of the radius variability in the experimental datasets. To conclude, cell length and cell volume can often be used interchangeably owing to small cell width fluctuations.

## 1 Introduction

Recent developments in microscopy and microfluidics have enabled researchers to study cell cycle regulation at a single-cell level [1]. Data analysis methods and quantitative models complement these experiments allowing us to make progress in our understanding of cell size homeostasis. Coarse-grained phenomenological models agnostic of the molecular details have provided clues about cell size regulation mechanisms [2–4]. Often, the Pearson correlation coefficient and best linear fit predictions from models are compared against experimental data to uncover the underlying biological mechanism. We show an example (Figure 1A) where the adder model of cell division - cells divide upon adding a constant size from birth (black dashed line) - is consistent with the experimental data (red points) of *Escherichia coli* [5–9]. In these analyses, cell length is assumed to be a proxy for cell volume and they are often used interchangeably.

**Figure 1:**
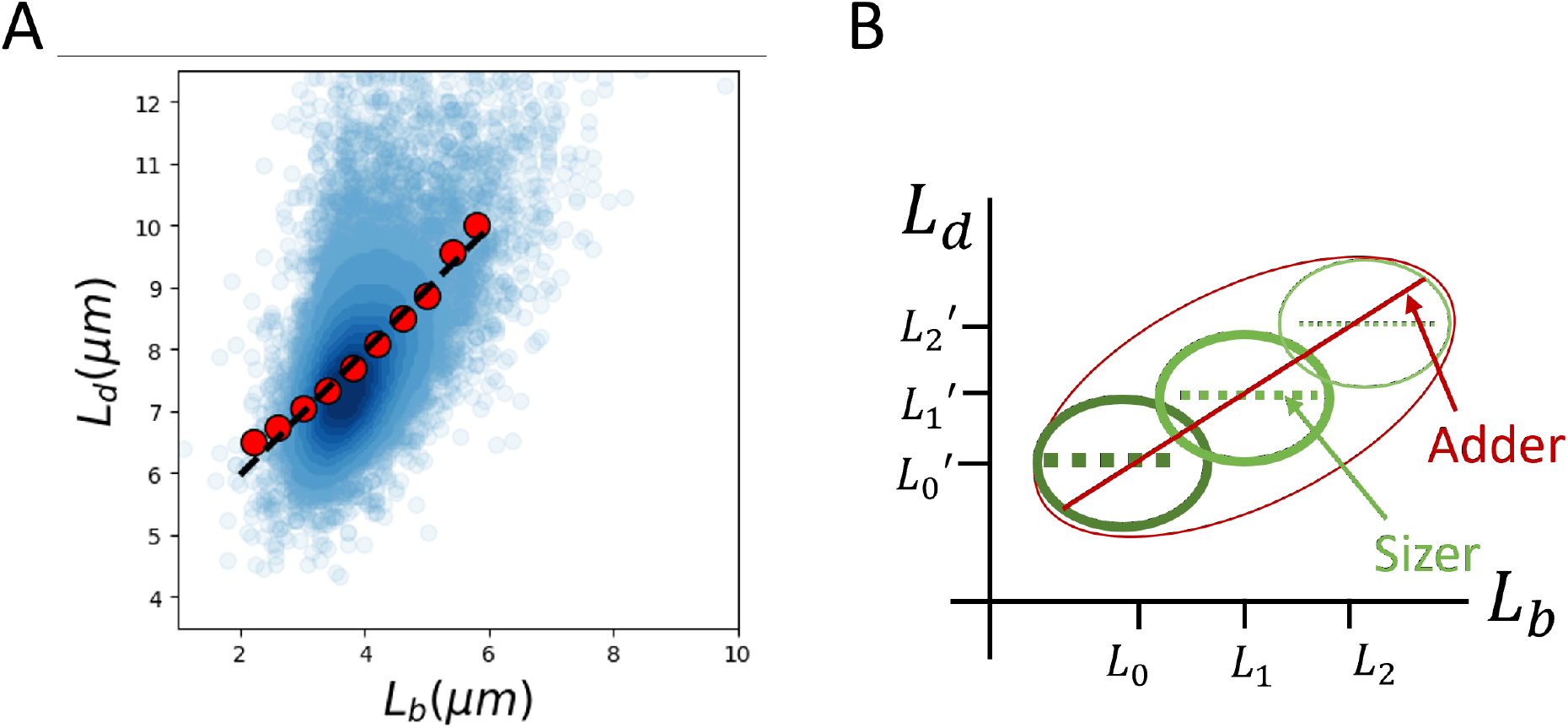
**A**. An example where the adder model of cell division, in which cells divide upon adding a constant size from birth is consistent with the experimental data of *E. coli*. The blue cloud is the raw data obtained from Ref. [6] growing in a growth medium with the average doubling time (⟨*T*_*d*_⟩) = 17 min (no. of cells N = 18248). The experimental binned data trend (red dots) matches the prediction for the binned data trend of the adder model (black dashed line with slope = 1).**B**. Illustration of Simpson’s paradox affecting length correlations. The underlying model could be a volume sizer with width fluctuations. Each green cloud corresponds to cells of different radii and is a volume sizer. Each subpopulation with a different radius is also a length sizer but with different average birth and division lengths. When the population of cells of different radii is combined, the correlation resembles that of a length adder.

Ref. [10] argued that they are are not interchangeable cell characteristics. Their logic followed a scenario pointed out in Ref. [2]. Ref. [10] discussed that subpopulations with different mean lengths at birth and division can arise despite carefully maintained growth conditions in microfluidics experiments due to natural cell width variability. For a population of cells with multiple subpopulations differing in their average length at birth and division, a division strategy called the sizer model, where cells divide upon reaching a critical size (slope = 0 in *L*_*d*_ vs *L*_*b*_ plot in each subpopulation) can be misconstrued as an adder (slope = 1 in *L*_*d*_ vs *L*_*b*_ plot) when the whole population is considered (Figure 1B). This is the so-called “Simpson’s paradox” and in this case leads to the actual sizer model being interpreted as an adder when various subpopulations are analyzed together. They found that cell width fluctuations could explain the positive length correlations observed in experimental data on fission yeast despite each subpopulation being a sizer. Using the same argument for *E. coli*, they found that a volume sizer along with cell width fluctuations could lead to a slope = 1 for the best linear fit of *L*_*d*_ vs *L*_*b*_ plot. The underlying biological mechanisms leading to a sizer differ from those resulting in an adder. This would mean that cell cycle analyses involving cell length data requires width fluctuations to be accounted for.

In this paper, we discuss various cell division models that take cell width fluctuations into account. First, we show that the model proposed in Ref. [10] is inconsistent with experimental data from *E. coli* regarding the correlation between birth lengths in successive generations. Based on a general model of cell division and cell width fluctuations, we devise a method involving conditional correlation of length variables conditioned upon radius variables to elucidate the underlying division volume control mechanism. Upon accounting for the width fluctuations, we find *E. coli* experimental data to be consistent with an adder model of cell division. Further, we estimate the contribution of measurement noise to the measured radius variability and conclude that intrinsic (actual) cell width fluctuations are smaller than those observed experimentally. This supports the assumptions from previous studies that cell lengths and volumes can be used interchangeably.

## 2 Results and Discussion

First, we will state the model proposed in Ref. [10] and obtain the correlation structure based on it. Using the model, we will restate the results in Ref. [10] and Figure 1B related to the correlation between the length at birth and division. Next, we will show that the minimal model fails to recover the correlations between birth lengths in successive generations.

In the model, *E. coli* single-cell geometry is approximated as a spherocylinder with identical cell radii (*R*) within a cell lineage (Figure 2A). However, in a typical cell cycle analysis such as the one in Figure 1A, the data combines cells from different lineages which might have different cell widths (and equivalently radii). One scenario where such cell radii variation might arise is if the cell width varies at a timescale much larger than the doubling time of cells. In the subsequent section, we will discuss a model where the cell radius changes in consecutive generations of a cell lineage. We assume that the cell divides when it reaches a critical volume 2*V*_0_ (volume sizer). This is a special case of the general model where the volume at birth determines the cell division volume via a regulation strategy, *f* (*V*_*b*_) [11]. A simple choice of *f* (*V*_*b*_) is *f* (*V*_*b*_) = 2(1−*α*)*V*_*b*_ +2*αV*_0_, where *α* is the cell cycle regulation parameter. Mathematically, we can express the division volume to be *V*_*d*_ = 2(1−*α*)*V*_*b*_ +2*αV*_0_(1+*ζ*_*s*_), where *ζ*_*s*_ is the size additive division noise. *α* can assume any value between 0 and 2 with *α* 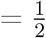 being the adder strategy, and *α* = 1 the sizer strategy. The cells divide symmetrically on average i.e., the mean division ratio 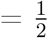 and the noise in division ratio for cells dividing in generation *n* = *δ*_*n*_. Upon dividing, the two daughter cells develop hemispherical poles at one end keeping the total volume constant i.e., the division volume of the mother cell is equal to the sum of birth volumes of the two daughter cells. This amounts to an addition of 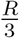 term to the birth lengths (“Pole formation after division” model in Figure 2A). Ref [10] based the model on fission yeast growth where a rapid increase in cell lengths is observed just after cell birth [12]. A similar new pole formation pattern is observed in bacterial species such as *Bacillus subtilis* [13]. In the next section, we will discuss a model of pole formation that is found in *E. coli*, where cells start constricting and form the new hemispherical poles before division.

**Figure 2:**
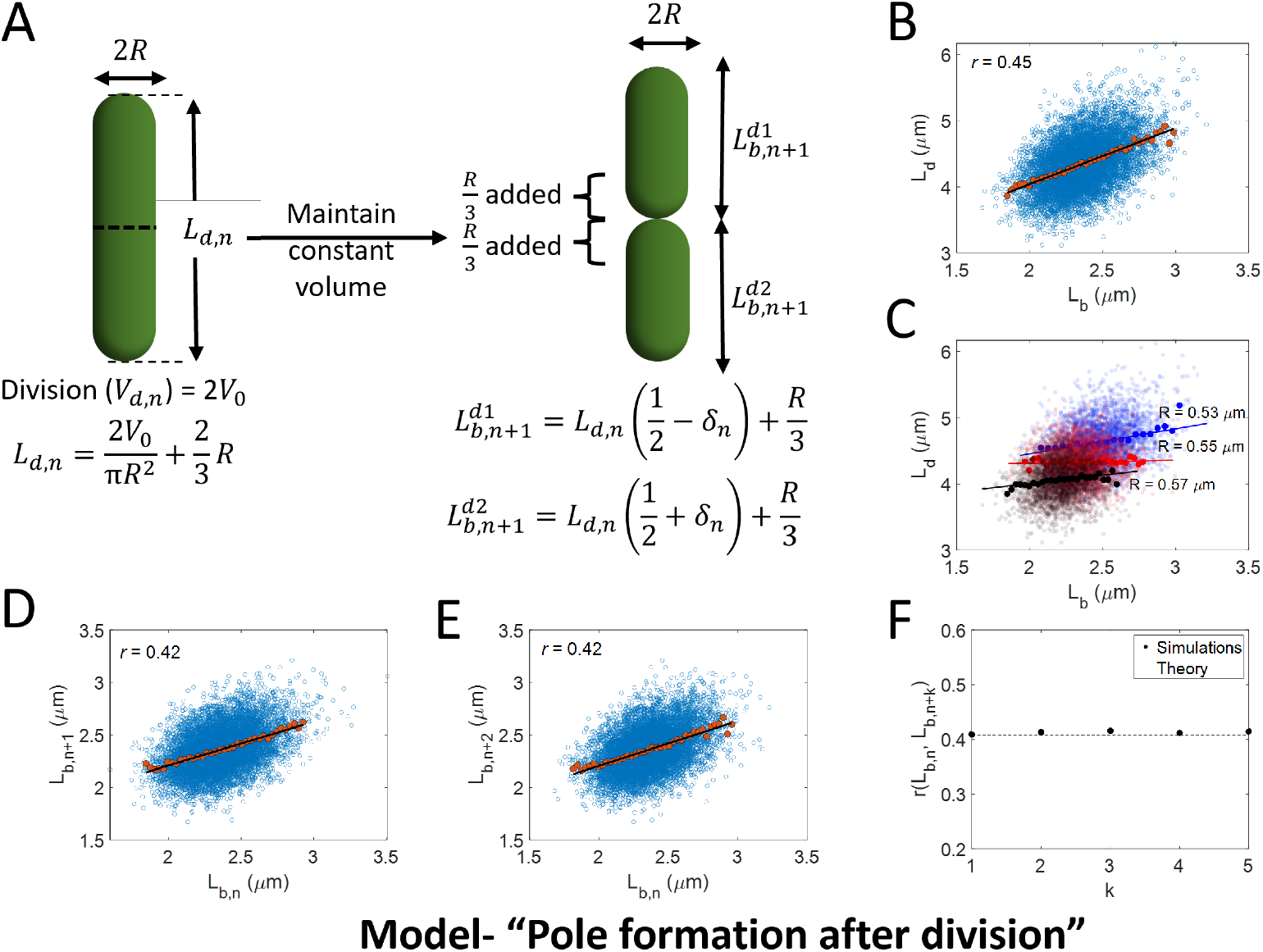
Model “Pole formation after division”. **A**. Schematic of the model proposed in Ref. [10]. The cell divides when it reaches a critical volume 2 *V*_0_ (volume sizer). The cells divide symmetrically by length on average i.e., the mean division ratio 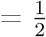 and the noise in division ratio for cells dividing in generation *n* = *δ*_*n*_. Upon dividing, the two daughter cells develop hemispherical poles at one end rapidly keeping the total volume constant i.e., the division volume of the mother cell is equal to the sum of birth volumes of the two daughter cells. This amounts to an addition of 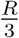 term to the birth lengths. The cell radius is fixed for all cells in a particular lineage but varies for different lineages. **B-E**. Simulations of the model in Figure 2A are carried out for 10000 cell lineages and over 25 generations. For the simulations, **B**. we plot *L*_*d*_ vs *L*_*b*_ plot. The correlation *r* (top left) points to a near-adder model. **C**. We plot *L*_*d*_ vs *L*_*b*_ for small subsets of *R*. We arrange the simulated dataset in ascending order and divide it into three groups. For each group (with a different average radius), we plot the *L*_*d*_ vs *L*_*b*_ plot. The plot shows the Simpson’s paradox mentioned in Figure 1B. **D**. Length at birth in generation *n* + 1 vs generation *n* is plotted. **E**. Length at birth in generation *n* + 2 vs generation *n* is plotted. The correlation values are identical and consistent with Eq. 1. In all the plots, the cloud is the raw data, the dots represent the binned data, and the line is the best linear fit. **F**. We calculate the correlation between the lengths at birth in generation *n* and *k* generations later for different *k*. Consistent with Eq. 1 and Figures 2D-2E, we find the correlations to be constant independent of *k*.

Upon simulating the model with the same model parameters as in Ref. [10], we recover their result of slope ≈ 1 for the best linear fit of *L*_*d*_ vs *L*_*b*_ plot which is consistent with a length adder (Figure 2B and Section S1.1.1 in SI text). As stated previously, this positive correlation comes from Simpson’s paradox (Figure 1B): Consider cells with radii *δR* larger than the average. Such cells would have an average length at birth and division different from that of cells with radii *δR* smaller than average. Within each subpopulation, cells divide upon reaching a critical volume or equivalently critical length as the cell width within the subpopulation is fixed (Figure 2C). On combining the different subpopulations, the shifts due to different length averages lead to an apparent length adder (Figure 2B).

Minimal cell division models that neglect width fluctuations are consistent with the experimental data regarding the correlation between cell birth lengths in successive generations. Ref. [4] analyzed experimental data on *E. coli* from Ref. [14] and found the Pearson correlation coefficient between the birth lengths of mother and daughter cells to be approximately 0.5. The correlation between the birth lengths of mother and granddaughter cells was approximately 0.25, in accordance with the models. We tested whether the model that includes width fluctuations also agrees with these correlations. Using the model, we found the correlation between length at birth at generation *n* (*L*_*b,n*_) and k generations later (*L*_*b,n*+*k*_) (Section S1.1.2 in SI text) to be,

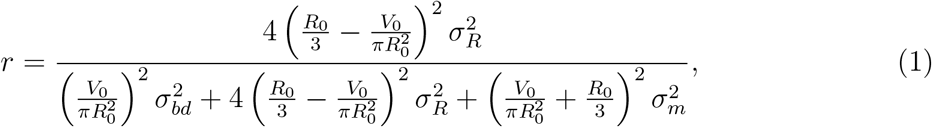

where *R*_0_ is the mean radius, *σ*_*bd*_ is the standard deviation of division noise *ζ*_*s*_, *σ*_*R*_ is the coefficient of variation (CV) in radius, and *σ*_*m*_ is the standard deviation of division ratio *δ*_*n*_. Contrary to the decreasing correlations with increasing *k* that have been observed for experimental data [4], we found that correlations between successive birth lengths are a constant and independent of *k* for this model (Eq. 1 and Figures 2D-2F). In this model, cell lengths have no memory of the prior generations (sizer) except for having the same radius. Since the radius stays constant within a lineage independent of *k*, the correlations do not vary despite considering cell lengths *k* generations apart. This is evident in Eq. 1 where the covariance between birth lengths *k* generations apart (the numerator in Eq. 1) is only dependent on the variability in cell width. The positive correlations in this case have the same origin as before i.e., Simpson’s paradox (Figure S1).

Thus, our analysis shows that a volume sizer with width fluctuations cannot account for the correlations between birth lengths observed in experimental data.

### Correlation structure in a cell division model where the new pole forms at mid-cell before division

In the previous model, hemispherical pole formation followed the cell division event. We wanted to verify that the timing of the new pole formation does not affect the correlation structure obtained from the model. The model in the previous section assumed a sudden increase in cell length in a small period around birth which is not observed in *E. coli*. Therefore, we test a model of cell growth and division where cells constrict and form new poles before dividing (“Pole formation before division” model).

The model borrows characteristics from the previous one, such as spherocylindrical cell geometry, constant radius along a lineage while variability exists across lineages, and a volume sizer. However, in this model, we assume that two spherocylindrical daughter cells are fully formed just before division. The two daughter cells are symmetrical on average but there are fluctuations in the volume division ratio denoted by *δ*_*n*_. For an asymmetrical division in generation *n*, it is assumed that one of the cells receives an additional *δ*_*n*_*V*_*d,n*_ in volume. However, the length partition for that cell is not *δ*_*n*_*L*_*d,n*_ since cell length is a linear function of volume (Figure S2A). Partitioning by length does not impact the results qualitatively (Figures S3A-S3C).

We find that, for the same parameters as in the previous section, the Pearson correlation coefficient between *L*_*b*_ and *L*_*d*_ is close to 0.5, thus, agreeing with a length adder (Figure S2B). It follows the same explanation as the previous section where Simpson’s paradox leads to an apparent adder (Figure S2C). Importantly, the correlation between birth lengths, *k* generations apart, is a constant for *k*=1 and 2 (Figures S2D-S2E and Section S1.2 in SI text). As was the case previously, the constancy in correlation *k* generations apart is because the lengths are correlated owing to the same radius in a lineage. Thus, the model fails in a similar manner as the previous model.

### Correlation structure in a model where radius correlations between generations are not one

Both the previous models show that volume sizer can lead to a length adder but they do not agree with all the correlations observed in the experiments. The correlations between length births across generations do not decay because the radius in a lineage is fixed. However, if the cell width fluctuates on a timescale of cell doubling time, then, the radius might be correlated but not equal to one between successive generations. In this section, we will discuss a model where we relax the assumption that the radius is fixed in a lineage (“Changing radii in a lineage” model).

The model discussed here (Figure 3A) varies in two aspects from the model discussed in the previous section. First, we consider a general model of cell division where the division volume *V*_*d*_ is determined via the regulation strategy *f* (*V*_*b*_) discussed previously. Further, we relax the assumption that the cell width is fixed for a few generations in a lineage. During the cell cycle, there could be changes to the cell width such that the radius just before the cell division in generation *n* is,

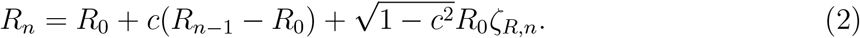

The radius is correlated with that in the previous generation (*R*_*n*−1_), *c* being the Pearson correlation coefficient of radii between generations. *ζ*_*R*_ is the noise in radius assumed to be Gaussian with mean 0 and standard deviation *σ*_*R*_. We assume that the radius does not change during the cell separation following the completion of septum formation. Thus, there are no sudden changes in cell lengths following the cell division process. The model also incorporates the spherocylinder cellular geometry and the two fully formed daughter cells at division features from the “Pole formation before division” model.

**Figure 3:**
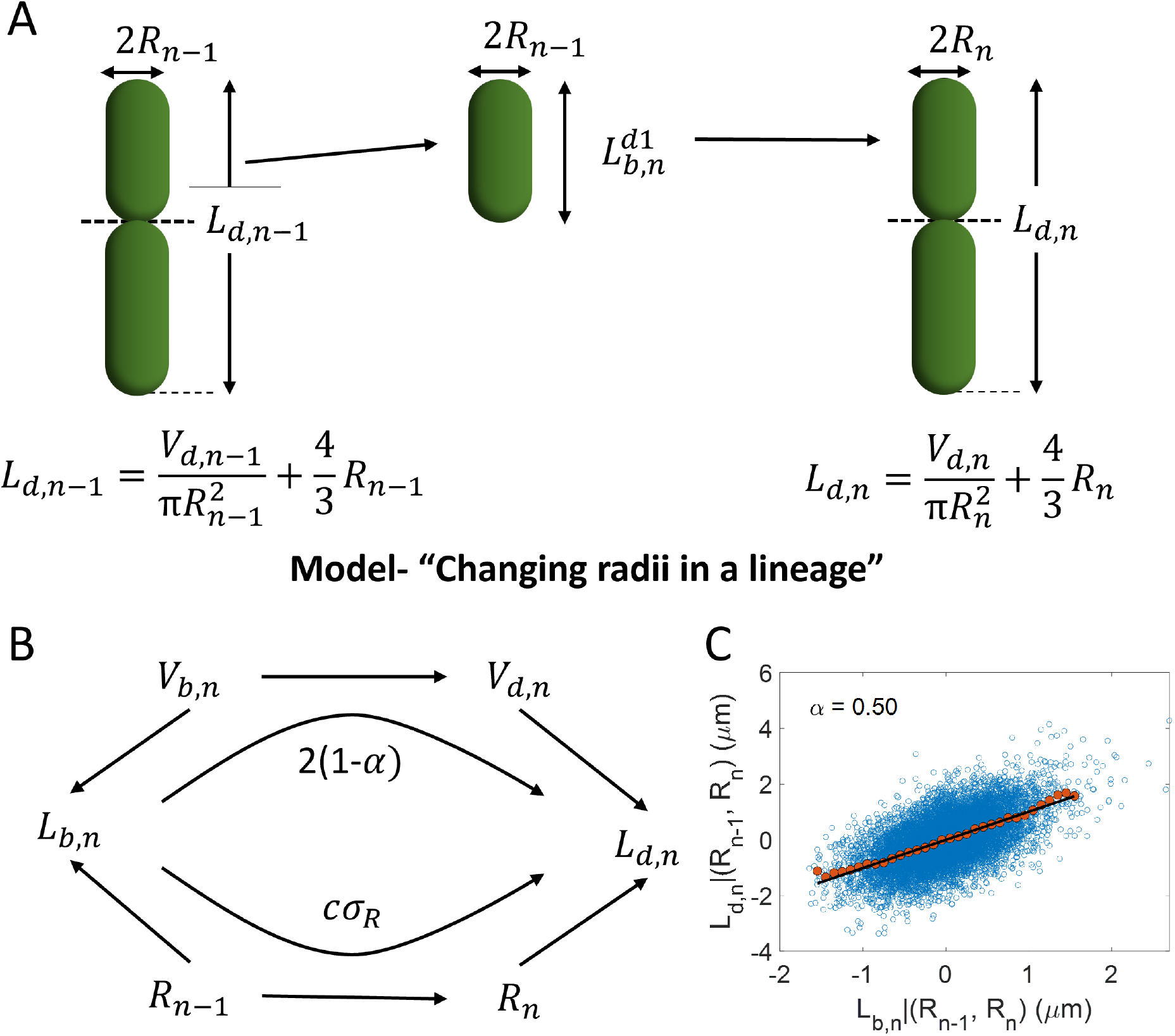
Model “Changing radii in a lineage”. **A**. Schematic of the model. The model assumes that cells have a spherocylindrical geometry and they divide based on their birth size. There are already two fully formed cells just before cell division. The cells divide symmetrically by volume on average. The cell radii are correlated but not the same for successive generations. **B**. Causal diagram showing that length at birth and division are correlated via two paths. One contribution is from the correlated radii in successive generations and other is from the division volume regulation strategy. The arrows in the graph point from cause to effect. **C**. Residuals *L*_*d,n*_|(*R*_*n*−1_, *R*_*n*_) vs *L*_*b,n*_|(*R*_*n*−1_, *R*_*n*_) are plotted for simulations of the model in Fig. 3A with *α* = 0.5. Using the slope of the best linear fit of the plot, we obtain an *α* (top left) consistent with the *α* used in the simulations.

Similar to the previous section, we can calculate the correlation between birth lengths *k* generations apart,

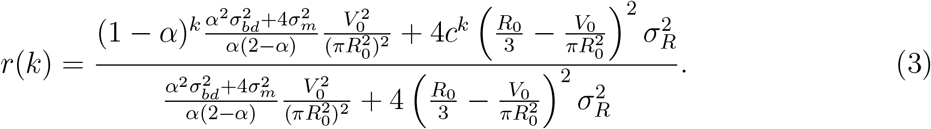

The length at birth in generation *n* + 1 (or equivalently the length at division in generation *n*) is related to the birth length in generation *n* through two paths (Figure 3B) - 1. via the cell cycle regulation strategy, 2. through the correlated cell radius across generations. The regulation strategy *f* (*V*_b_) leads to a correlation contribution of 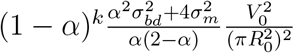 between birth volumes *k* generations apart (first term in Eq. 3). The 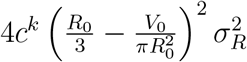 term in Eq. 3 is the contribution of correlated radii across generations to the birth length correlations. We validate Eq. 3 using simulations of the “Changing radii in a lineage” model. The correlations between birth lengths *k* generations apart were consistent between simulations and theory in Eq. 3 (Figure S4A). Further, theory and simulation results for the correlation between birth lengths were consistent for varying cell width variability, *σ*_*R*_ (Figure S4B). If 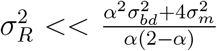, we recover the (1 − *α*)^*k*^ decay of correlations with increasing *k*, consistent with a regulation strategy *f* (*V*_*b*_) and neglecting radius fluctuations (Figure S4B).

### Determining the volume cell cycle regulation strategy

We calculated the expression for the correlation between birth lengths *k* generations apart based on a general model of cell division which accounts for cell width fluctuations. The cell width fluctuations confound the length correlations, thus, hiding the value of the cell cycle regulation parameter, *α*. In this section, we discuss a method based on conditional correlations to estimate the value of *α* provided that the length at birth, the division length, and the cell radius for mother and daughter cells are measured.

A recent study used conditional correlations to restrict the cell cycle model space [15]. The method entails calculating the correlation between two variables upon fixing the value of the third variable(s). This is equivalent to finding how two variables are correlated given the effects of the third variable(s) are removed. On removing the contribution of cell radius to cell lengths, a correlation between the transformed cell lengths would solely arise from the cell regulation strategy path illustrated in Figure 3B. The length variables *L*_*b,n*_ and *L*_*d,n*_ are dependent on radius values *R*_*n*−1_ and *R*_*n*_ respectively in the general model discussed previously (Figure 3A). The residuals obtained upon linear regression of length variables, *L*_*b/d,n*_, on both radius *R*_*n*−1_ and *R*_*n*_ (*L*_*b/d,n*_|(*R*_*n*−1_, *R*_*n*_)) denotes variables which have no contribution from cell radius. The slope of the best linear fit of *L*_*d*_|(*R*_*n*−1_, *R*_*n*_) vs *L*_*b*_|(*R*_*n*−1_, *R*_*n*_) plot is equal to 2(1 − *α*) (Section S1.2.2 in SI text). We verified it using simulations where we fixed the value of *α* to be 0.5 (Figure 3C). The *α* estimate obtained using the conditional correlation has only small errors when measurement errors in cell width measurements and correlated measurement noise are included (Section S1.2.2 in SI text and Figure S5). Ref. [10] use a similar method where fission yeast cells were pooled into subsets based on their radii. They find that the slope of the best linear fit of the *L*_*d*_ vs *L*_*b*_ plot tends to zero upon decreasing the variability in radius, thus, agreeing with a sizer (*α* = 1).

### Test on experimental data

Next, we used the conditional correlation method on experimental datasets of *E. coli* collected in Ref. [6]. Using microfluidic devices called mother machines, the study measures the birth lengths, division lengths, and the cell radius for multiple single cells. Upon carrying out the conditional correlation analysis, we find the correlation to be close to 0.5 (Table 1). We show the plot for one of the growth conditions in Figure 4A. Thus, the *α* ≈ 0.5 observed from experimental data implies that the underlying cell size regulation strategy cannot be a volume sizer. Similar estimates of the conditional correlation (*L*_*b,n*_, *L*_*d,n*_)|(*R*_*n*−1_, *R*_*n*_) was obtained using *E. coli* data in other studies (Table S1 [16], Table S2 [9]).

**Table 1:** Pearson correlation coefficients along with their 95% confidence intervals (CI) are shown for *E. coli* cells growing in six different growth media in Ref. [6] with mean generation times, ⟨*T*_*d*_⟩. Correlation between *L*_*b*_ and *L*_*d*_, *L*_*b,n*_ and *L*_*b,n*+1_ i.e., *r*(1), and conditional correlation, (*L*_*b,n*_, *L*_*d,n*_)|(*R*_*n*−1_, *R*_*n*_), are shown.

| $\langle T_d \rangle$<br>(min) | No. of cells | $(L_{b,n}, L_{d,n})$ | $(L_{b,n}, L_{d,n})\vert (R_{n-1}, R_n)$ | $r(1); (L_{b,n}, L_{b,n+1})$ |
| --- | --- | --- | --- | --- |
| 17 | 17055 | 0.55 (0.54,0.56) | 0.57 (0.56,0.58) | 0.52 (0.51, 0.53) |
| 22 | 16636 | 0.49 (0.48,0.50) | 0.49 (0.48,0.51) | 0.49 (0.48, 0.50) |
| 27 | 17682 | 0.56 (0.55,0.57) | 0.56 (0.55,0.57) | 0.53 (0.52, 0.54) |
| 31 | 28808 | 0.53 (0.52,0.54) | 0.54 (0.53,0.54) | 0.53 (0.52, 0.53) |
| 39 | 20300 | 0.56 (0.55,0.56) | 0.55 (0.55,0.56) | 0.56 (0.55, 0.57) |
| 51 | 9763 | 0.55 (0.54,0.57) | 0.54 (0.52,0.55) | 0.59 (0.57, 0.60) |

**Figure 4:**
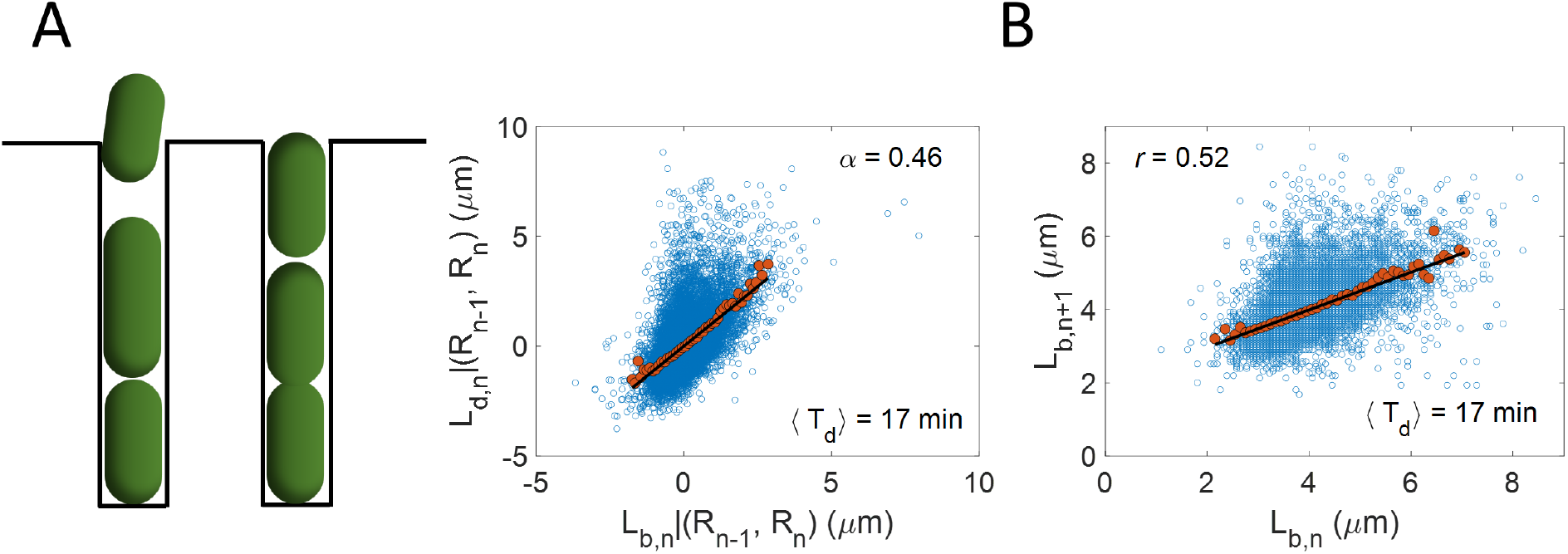
**A**. Schematic diagram of a mother-machine (left). Residuals *L*_*d,n*_ |(*R*_*n*−1_, *R*_*n*_) vs *L*_*b,n*_ |(*R*_*n*−1_, *R*_*n*_) are plotted (right) for *E. coli* experimental data obtained using mother-machine experiments in Ref. [6] (⟨*T*_*d*_⟩ = 17 min, no. of cells = 17055). The *α* value calculated is noted in the top left. **B**. *L*_*b,n*+1_ vs *L*_*b,n*_ plot is made for the same dataset in Fig. 4A (see Section S3.1 in SI text for details about data analysis).

### Implications on cell width fluctuations

We can use the “Changing radii in a lineage” model to predict the correlation between birth lengths in successive generations (*k* = 1 in Eq .3) Substituting the value of *α*, obtained using the slope of the best linear fit of *L*_*d*_|(*R*_*n*−1_, *R*_*n*_) vs *L*_*b*_|(*R*_*n*−1_, *R*_*n*_) plot, along with other model parameters (determined using experiments as shown in Section S2.2 in SI text), into Eq. 3, we expect the correlation between birth lengths in consecutive generations to be 0.62-0.82 (Table S3). This is inconsistent with experimental results (Table 1 and Figure 4B) where the correlations are close to 0.5.

These estimates for correlation are based on the assumption that there is negligible measurement noise. However, the length and width measurements will have variability contributions from the intrinsic stochasticity of the biochemical reactions as well as from inaccuracies in length measurements. Cell widths, which are of the order of 200-500 nm, will have significant measurement errors owing to their small absolute magnitudes. Thus, the values of *c* and *σ*_*R*_ that are substituted into Eq. 3 will be imprecise. Next, we try to estimate the values of *c* and *σ*_*R*_ that might reconcile *α* = 0.5 estimated previously with correlation = 0.5 from experiments.

We simplify Eq. 3 by assuming that the length of the cell is generally larger than the cell radius, i.e., 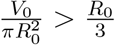. Thus, the correlation between birth lengths in consecutive generations is approximately only a function of noise variables and *c*. Substituting *α* =0.5 in Eq. 3,

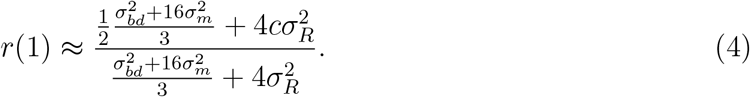

We find that *r*(1) will be 0.5 in two cases - either *c* = 0.5 or 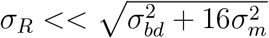. While either of these cases is possible in principle, strategies to regulate the radius across generations to be 0.5 are unknown. Thus, we conclude that the intrinsic variability of radius (*σ*_*r*_) is much smaller than the noise in setting division length. Most of the measured cell width variability has large contributions from measurement errors.

Until now, we have focused on models where cell volume is the relevant quantity the cell controls. The underlying assumption is that the rate of volume growth is proportional to the rate of protein synthesis and biomass accumulation. Ref [17] observed that the cell surface area to biomass ratio remains constant during the cell cycle while density changes. It would mean that cell surface area instead of cell volume is the appropriate proxy for biomass accumulation. However, our conclusions about the actual cell width variability being small remain unchanged even if we consider surface area being controlled (Section S4 in SI text).

A striking observation in the experimental data is that the correlations between *L*_*b,n*_ and *L*_*d,n*_ closely agree with the conditional correlations (*L*_*b,n*_, *L*_*d,n*_)|(*R*_*n*−1_, *R*_*n*_) (Table 1, Table S1 and Table S2). Next, we use this observation and cell cycle model simulations to restrict the value of *σ*_*R*_ and *c*. In the simulations, we allow for a difference of 0.02 between the two correlation values. Most of the correlation differences obtained from experiments are within this limit (Table 1, Table S1, and Table S2). We vary *σ*_*R*_ and *c* keeping the measured values of radius CV (*σ*_*Rt*_) and correlation between consecutive generations (*c*_*t*_) fixed. We restrict the parameter space in the simulations such that the correlation between measurement noise in radius in successive generations is between 0 and 1. For the division regulation strategy *f* (*V*_*b*_), we find that the correlation and conditional correlation are close for smaller values of *σ*_*R*_ (Figure S6A). We find that the measurement noise in width accounts for at least half of the total width variability. Similar results are obtained for cell cycle models beyond the *f* (*V*_*b*_) regulation strategy (Figures S6B-S6D).

Thus, the correlation and conditional correlation values bound the intrinsic cell width variability at most half the width variability observed in experiments.

## 3 Conclusion

Our previous work has shown that unaccounted sources of noise could lead to misinterpretation of the results of single-cell data analysis methods [18]. Ref. [10] presented another example of the misinterpretation where width fluctuations would lead to a volume sizer being misinterpreted as a length adder (Figure 1B and Figs. 2A-2B). In a volume sizer, division happens when a certain sizer protein reaches a threshold amount irrespective of its initial amount at birth, and the protein biosynthesis is coupled to volume growth. Note that the underlying mechanism might not involve sensing the absolute protein numbers in the cell. The sizer proteins can be limited to a region of particular size. The rise in their concentration or equivalently number in this region to a threshold amount can trigger division as is found in fission yeast [19]. In contrast, for a length adder, division happens when the protein accumulates a critical amount from birth and the protein biosynthesis is coupled to length growth. Thus, two differences arise between these two models-1. What controls division? - the protein accumulates by a threshold amount or the absolute number reaches a critical amount, 2. Is the biomass accumulation related to length or volume?

Cell volume is also a suitable candidate to be the proxy for cell size. Biochemical reaction kinetics in the cell depend on the concentration of reactant species which is tied to the cell volume. Ref. [20] also showed that the average cell volume rather than cell length, surface area, and width, scaled as *e*^⟨*λ*⟩(*C*+*D*)^, where ⟨*λ*⟩ is the average growth rate and *C* + *D* is the average time between the start of DNA replication and division. This observation can be explained by assuming that the average volume per origin of replication at the initiation of DNA replication is a constant and division happens after a constant *C* + *D* time from birth [21]. In fast and intermediate growth conditions, the *C* + *D* time was found to be 60 min [22]. Since cell volume is the only cell geometry characteristic that follows the growth law, it is natural to assume that cell volume per origin is regulated at the start of DNA replication [23]. Thus, assuming a volumetric control, we find from Eq. 4 that length and volume correlations can be the same if intrinsic radius variability is small compared to other fluctuations (division size, division ratio). Note that for a spherocylindrical cell geometry with negligible cell width variability, volume at birth or division is a linear function and not proportional to the cell length. However, cell cycle analysis frequently uses Pearson correlation coefficients which are invariant under linear transformations. Hence, it is often equivalent to use volume or length in the cell cycle data analysis.

The mechanistic understanding of width control in bacteria is still an open question of research [24–27]. Our results suggest that width control in *E. coli* is even more accurate than previously thought. We found that length correlations between birth and division, and the conditional correlation between them when conditioned upon radius in the mother cell and current generation was extremely close in experimental data in multiple growth conditions from multiple studies (Table 1, Table S1, and Table S2) [6, 9, 16]. Using simulations of cell cycle models (Figure S6), we found the range of intrinsic radius variability for which the above equality in correlations holds. The simulations indicate that the variance contribution of the measurement noise in radius can be at least half of the total radius variability. If the CV of the radius is approximately 6% (Table 1), then our estimates put the CV of the intrinsic radius variability to be less than 4% (standard deviation ≈ 10 nm). In general, we conclude that the measured cell width variability is dominated by measurement noise. Hence, more accurate measurements, possibly using electron microscopy, are required to measure the actual (intrinsic) radius variability.

More broadly, adder behavior has been observed in multiple species across different domains of life [2, 7, 28–30]. For a volume sizer to appear as a length adder requires fine-tuning of cell cycle parameters such as cell radius variability and noise in setting the division size. It is unlikely that the parameters are precisely controlled in these different organisms with different geometries and underlying cell cycle mechanisms. Why the adder behavior is so ubiquitous remains an open question.

## Author Contributions

A.A. and P.K. conceptualized the project, P.K. carried out the analysis, P.K. and A.A. wrote the draft and reviewed and edited the manuscript. A.A. acquired funding.

## Acknowledgments

We thank Martin Howard, Sven van Teeffelen, and Sattar Taheri-Araghi for the useful feedback on the manuscript. A.A. and P.K. acknowledge support from NSF CAREER 1752024. A.A. thanks the generous support of the Clore Center for Biological Physics and ERC-CoG 2023 101125981.

## Notes

### Competing Interest Statement

The authors have declared no competing interest.

