## Supplementary text for "Are cell length and volume interchangeable in cell cycle analysis?"

#### Supplemental Figures

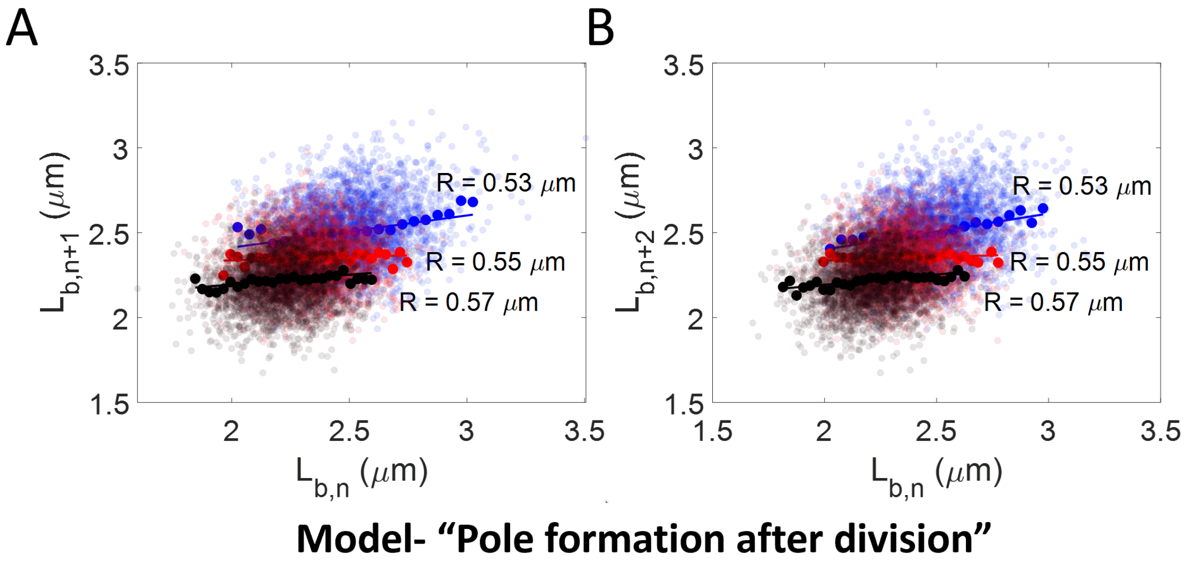


Figure S1: **Model "Pole formation after division" - A-B.** We simulate the model described in Figure 2 of the main text. We arrange the simulated dataset in ascending order of radius $R$ and divide it into three groups. For each group (with a different average radius), we plot the **A.** $L_{b,n+1}$ vs $L_{b,n}$ plot. **B.** $L_{b,n+2}$ vs $L_{b,n}$ plot. These plots illustrate Simpson's paradox.

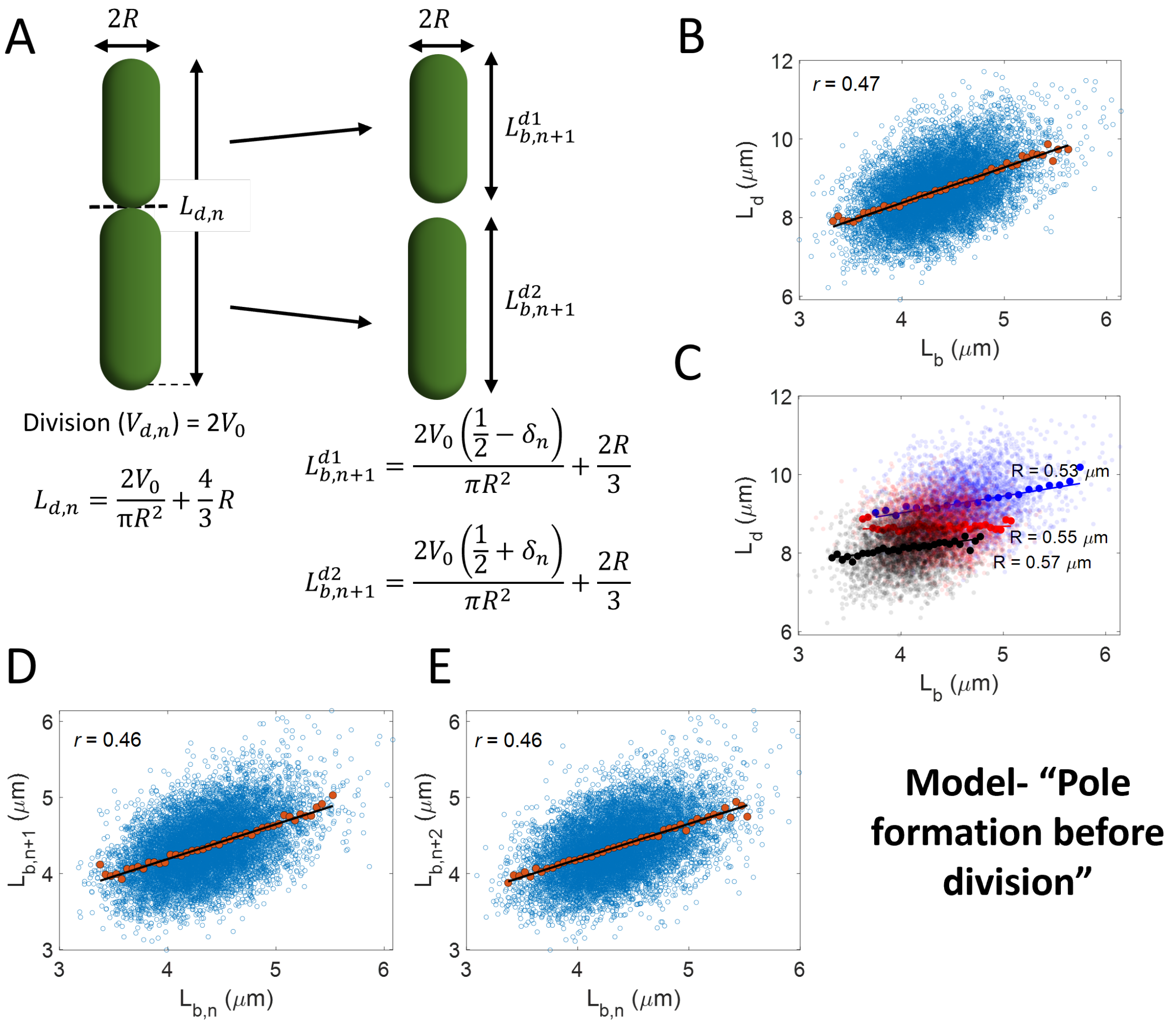


Figure S2: **Model "Pole formation before division" - A.** Schematic of the model proposed in Section "Correlation structure in a cell division model where the new pole forms at mid-cell before division" of the main text. The model borrows certain aspects from the model in Figure 2A of the main text. In both models, cells have a spherocylindrical geometry and they divide upon reaching a critical volume. The radius is fixed for a particular lineage and varies between lineages. However, unlike the model in Figure 2A, there are already two fully formed cells just before cell division. The cells divide symmetrically, on average, by volume. **B-E.** Simulations of the model in Figure S2A are carried out for 10000 cell lineages and over 25 generations. For the simulations, **B.** we plot $L_d$ vs $L_b$ plot. The correlation $r$ (top left) points to a near-adder model. **C.** We plot $L_d$ vs $L_b$ for small subsets of $R$. We arrange the simulated dataset in ascending order and divide it into three groups. For each group (with a different average radius), we plot the $L_d$ vs $L_b$ plot. The plot shows Simpson's paradox mentioned in Figure 1B. **D.** Length at birth in generation $n+1$ vs generation $n$ is plotted. **E.** Length at birth in generation $n+2$ vs generation $n$ is plotted. The correlation values are identical and consistent with Eq. S28. In all the plots, the cloud is the raw data, the dots represent the binned data, and the line is the best linear fit.

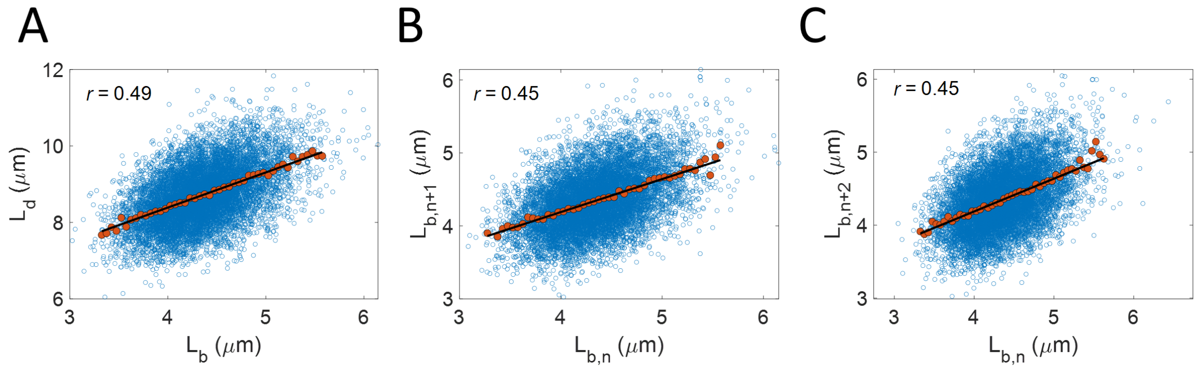


Figure S3: **A-C.** The simulation results presented here correspond to the cell cycle model in Figure S2A. The only difference between the two models is that cells divide symmetrically, on average, by length instead of volume. The length at birth for the daughter cell $L_{b,n+1} = L_{d,n}(\frac{1}{2} + \delta_n)$. Simulations of the model are carried out for 10000 cell lineages and over 25 generations. For the simulations, **A.** we plot $L_d$ vs $L_b$ plot. The correlation $r$ (top left) points to a near-adder model. **B.** Length at birth in generation $n+1$ vs generation $n$ is plotted. **C**. Length at birth in generation $n+2$ vs generation $n$ is plotted. The correlation values in all the plots are close to that in Figure S2. In all the plots, the cloud is the raw data, the dots represent the binned data, and the line is the best linear fit.

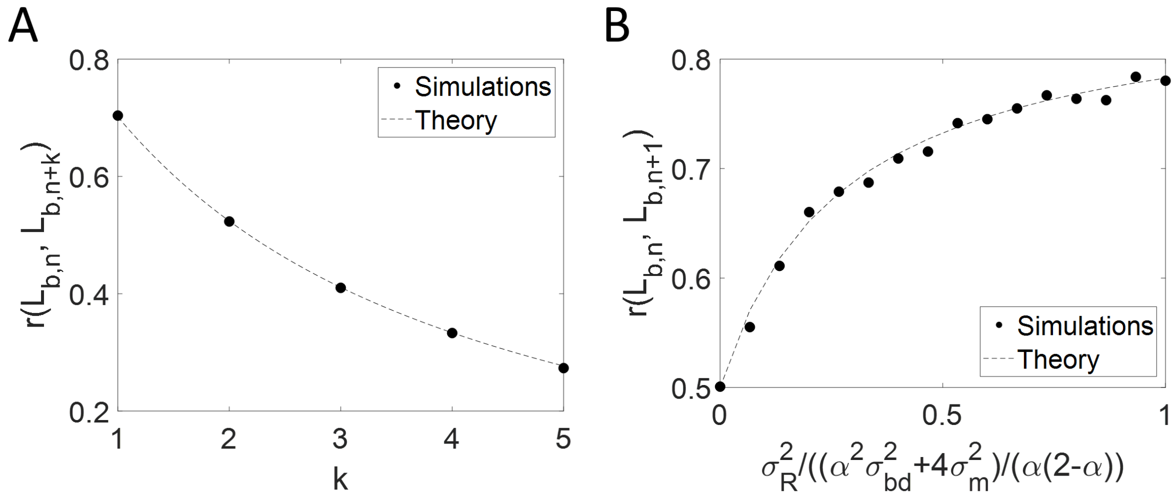


Figure S4: **Model "Changing radii in a lineage" - A-B.** We simulate the model described in Figure 3A of the main text. **A.** We find that the Pearson correlation coefficients between the lengths at birth in the $n^{th}$ generation ($L_{b,n}$) and $n+k^{th}$ generation ($L_{b,n+k}$) obtained from simulations (black dots) match the theoretical predictions (black dashed line) shown in Eq. 3 of the main text/Eq. S27 of the SI text. **B.** The coefficient of variation (CV) of the cell radius ($\sigma_R$) is varied in the simulations keeping the value of $\alpha = \frac{1}{2}$ (volume adder), size additive division size noise ($\sigma_{bd} = 0.19$, and noise in the division ratio ($\sigma_m = 0.03$) fixed. We find that the correlations between birth lengths in consecutive generations obtained from simulations (black dots) are consistent with the theoretical predictions (black dashed line) of Eq. 3 of the main text/Eq. S27 of the SI text for different values of $\sigma_R$.

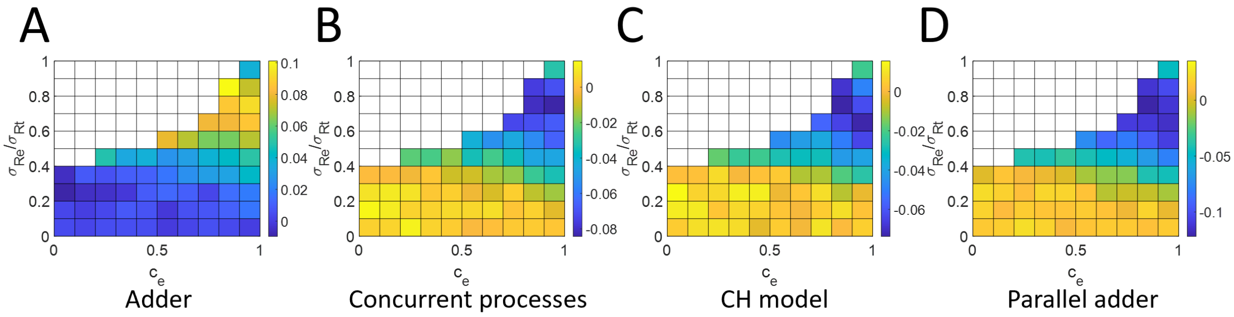


Figure S5: **Verifying the conditional correlation method to infer volume regulation strategies. A-D.** Differences between the correlation obtained from $V_d$ vs $V_b$ plot and $L_{d,n}|(R_{n-1}, R_n)$ vs $L_{b,n}|(R_{n-1}, R_n)$ plot are shown. In the section "Determining the volume cell cycle regulation strategy" of the main text, we put forth that the conditional correlation between the length at birth ($L_{b,n}$) and length at division ($L_{d,n}$) when conditioned upon radius in the current generation ($R_n$) and the previous generation $R_{n-1}$ is an appropriate method to study cell cycle regulation mechanisms despite width fluctuations. We simulate various cell cycle models with each plot representing a different model regulating the division size. In each of these simulations, we vary the measurement error in the cell radius measurements ($\sigma_{Re}$) and the correlation between the radius measurement errors in consecutive generations such that the correlation between the actual radius in consecutive generations is between 0 and 1. The white areas in the plots are the regions that do not follow this condition. The measured coefficient of variation (CV) of the radius ($\sigma_{Rt}$) and the measured correlations between the radius in consecutive generations ($c_t$) are fixed in the simulations. The differences in the actual and inferred volume regulation strategies are plotted. **A.** The division strategy is the adder model ($\alpha = \frac{1}{2}$). The values in the plot correspond to $\frac{1}{2} - \alpha_{infer}$ where $\alpha_{infer}$ is obtained using the slope of $L_{d,n}|(R_{n-1}, R_n)$ vs $L_{b,n}|(R_{n-1}, R_n)$ plot (Section S1.2.2). **B-D.** The plot values correspond to the difference between the Pearson correlation coefficients of $V_d$ vs $V_b$ plot and $L_{d,n}|(R_{n-1}, R_n)$ vs $L_{b,n}|(R_{n-1}, R_n)$ plot. **B.** The division strategy is the concurrent processes model [S1] where both birth and the start of DNA replication control the division volume (see S1.2.2 for details) **C-D.** The division volume is solely controlled by the volume at the start of DNA replication. **C.** Division happens after a constant time from the initiation of DNA replication [S2]. **D.** Division happens after an addition of constant volume per origin of replication from the initiation of DNA replication.

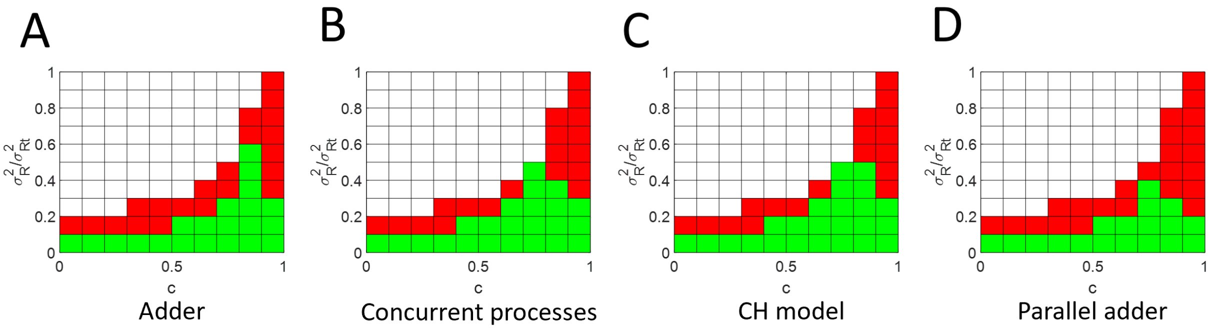


Figure S6: **A-D.** The differences between the conditional correlation $(L_{b,n}, L_{d,n}|(R_{n-1}, R_n)$ and correlation $(L_{b,n}, L_{d,n})$ are calculated. If the absolute magnitude of the difference is less than 0.02, the value is set to 1 (green) or else zero (red). The plots are made for simulations of the model in Fig. 3A of the main text. The difference between the four plots is the cell cycle model regulating the division size. The measured coefficient of variation (CV) of the radius $(\sigma_{Rt})$ and the measured correlations between the radius in consecutive generations $(c_t)$ are fixed in the simulations. The standard deviation of the actual radius fluctuations $(\sigma_R)$, and the correlation between the actual radii in consecutive generations $(c)$ is varied such that the correlation between the measurement noise in radius in consecutive generations $(c_e)$ is between 0 and 1. The white areas in the plots are the regions that do not follow this condition. **A.** Division volume is determined by the birth volume. **B.** The division strategy is the concurrent processes model [S1] where both birth and the start of DNA replication control the division volume (see S1.2.2 for details) **C-D.** The division volume is solely controlled by the volume at the start of DNA replication. **C.** Division happens after a constant time from the initiation of DNA replication [S2]. **D.** Division happens after the addition of constant volume per origin of replication from the initiation of DNA replication.

### Supplemental Tables

| $\langle \mathbf{T_d} \rangle$ (min) | No. of cells | $(\boldsymbol{L_{b,n}, L_{d,n}})$ | $(\boldsymbol{L_{b,n}, L_{d,n}})\vert(\boldsymbol{R_{n-1}.R_n})$ | $(\boldsymbol{L_{b,n}, L_{b,n+1}})$ |
|---|---|---|---|---|
| 74 | 298 | 0.50 (0.41, 0.58) | 0.48 (0.41, 0.58) | 0.34 (0.23 0.43) |
| 82 | 796 | 0.62 (0.57, 0.66) | 0.62 (0.58, 0.66) | 0.56 (0.51 0.61) |
| 110 | 430 | 0.56 (0.49, 0.62) | 0.54 (0.47, 0.61) | 0.49 (0.42 0.56) |

Table S1: Pearson correlation coefficients along with their 95% confidence intervals (CI) are shown for three datasets of untreated *E. coli* cells in Ref. [S3] with mean generation times, $\langle T_d \rangle$. Correlation between lengths at birth ($L_{b,n}$) and division ($L_{d,n}$), lengths at birth in consecutive generations ($L_{b,n}$ and $L_{b,n+1}$), and conditional correlation, $(L_{b,n}, L_{d,n})|(R_{n-1}, R_n)$, are shown. $R_n$ is the cell radius when the cell divides in generation $n$.

| $\langle \mathbf{T_d} \rangle$ (min) | No. of cells | $(\boldsymbol{L_{b,n}, L_{d,n}})$ | $(\boldsymbol{L_{b,n}, L_{d,n}})\vert(\boldsymbol{R_{n-1}.R_n})$ | $(\boldsymbol{L_{b,n}, L_{b,n+1}})$ |
|---|---|---|---|---|
| 29 | 757 | 0.55 (0.49, 0.59) | 0.54 (0.49, 0.59) | 0.54 (0.49 0.59) |
| 43 | 854 | 0.52 (0.47, 0.56) | 0.53 (0.47, 0.57) | 0.56 (0.52 0.61) |
| 52 | 1482 | 0.50 (0.47, 0.54) | 0.53 (0.49, 0.56) | 0.49 (0.45 0.53) |
| 64 | 992 | 0.51 (0.46, 0.55) | 0.54 (0.49, 0.58) | 0.44 (0.39 0.49) |
| 113 | 925 | 0.60 (0.56, 0.64) | 0.59 (0.55, 0.63) | 0.63 (0.59 0.67) |
| 194 | 680 | 0.49 (0.43, 0.54) | 0.48 (0.42, 0.54) | 0.44 (0.37 0.50) |

Table S2: Pearson correlation coefficients along with their 95% confidence intervals (CI) are shown for *E. coli* growing in six different steady-state growth conditions [S4] with mean generation times, $\langle T_d \rangle$. Correlation between lengths at birth ($L_b$) and division ($L_d$), lengths at birth in consecutive generations ($L_{b,n}$ and $L_{b,n+1}$), and conditional correlation, $(L_{b,n}, L_{d,n})|(R_{n-1}, R_n)$, are shown.

| $\langle \mathbf{T_d} \rangle$ (min) | $\mathbf{V_0}$ $\mu\mathbf{m^3}$ | $\sigma_{\mathbf{bd}}$ | $\sigma_{\mathbf{m}}$ | $\mathbf{R_0}$ $\mu\mathbf{m}$ | $\sigma_{\mathbf{R}}$ | **c** | $\alpha$ | **r(1) - Eq. 3, main text** |
|---|---|---|---|---|---|---|---|---|
| 17 | 2.77 | 0.19 | 0.03 | 0.49 | 0.08 | 0.86 | 0.46 | 0.72 |
| 22 | 1.70 | 0.17 | 0.03 | 0.42 | 0.07 | 0.85 | 0.55 | 0.67 |
| 27 | 1.04 | 0.09 | 0.02 | 0.35 | 0.05 | 0.84 | 0.50 | 0.73 |
| 31 | 0.77 | 0.04 | 0.02 | 0.34 | 0.07 | 0.90 | 0.52 | 0.82 |
| 39 | 0.58 | 0.10 | 0.02 | 0.32 | 0.06 | 0.91 | 0.52 | 0.76 |
| 51 | 0.46 | 0.14 | 0.04 | 0.27 | 0.06 | 0.84 | 0.57 | 0.62 |

Table S3: Model parameter values substituted to Eq. 3 of the main text and $r(1)$ values obtained from it. The model parameters are obtained using experimental data from Ref. [S5] and explained in SI text.

### S1 Rejecting models where volume sizer leads to length adder

#### S1.1 Volume sizer model in Ref. [S6]

Previous works [S6, S7] have shown that unaccounted noise variables can lead to misinterpretations of regression and correlation values. The main text focuses on one such example [S6] where the cell width fluctuations cause the actual underlying mechanism of a volume sizer (cell divides upon reaching a critical volume) to appear as a length adder (division happens upon constant length increment from birth). We will demonstrate it in this section by analytically calculating the expressions for the correlations between various cell cycle variables (such as length at birth and length at division) using simple cell cycle regulation models. Note that some of the calculations are shown here to introduce the model and motivate the question but these results are also presented in Refs. [S8], [S6], and [S7]. Further, we will show model correlation predictions that do not agree with the experimental data.

We use a simplistic model of cell division that was proposed in Ref. [S9]. In the model, cells divide when they reach a volume $V_d$ determined solely by their birth volume, $V_b$. Mathematically, we model this size regulation strategy as $V_d = f(V_b)$, where $f(V_b) = 2(1-\alpha)V_b + 2\alpha V_0$. $\alpha$ denotes the strength of the size regulation strategy and $V_0$ is the average cell volume at birth. $\alpha = \frac{1}{2}$ corresponds to the above-mentioned adder strategy while $\alpha = 1$ points to the sizer strategy. To account for the stochasticity in the cell division process, we introduce a size additive noise in addition to $f(V_b)$,

$$V_d = 2(1-\alpha)V_b + 2\alpha V_0(1 + \zeta_s(0, \sigma_{bd})). \tag{S1}$$

In the simulations, the noise $\zeta_s(0, \sigma_{bd})$ is drawn from a normal distribution with zero mean and standard deviation of $\sigma_{bd}$. $\zeta_s$ in successive generations are independent of each other.

Note that we use a size-additive noise due to ease of calculations, however, the nature of division noise (time or size additive) does not affect the results qualitatively [S7].

Assuming a spherocylinder geometry for the rod-shaped *E. coli* cells, we find the length at division ($L_d$) to be,

$$L_d = \frac{V_d}{\pi R^2} + \frac{2}{3}R, \tag{S2}$$

where $R$ is the cell radius. The cell radius is assumed to fluctuate on timescales greater than the doubling time such that cells within a few generations of a particular lineage will have the same radius while those from different lineages (with common ancestors multiple generations ago) will have different radii. The radius for different lineages is drawn from a normal distribution with $R_0$ mean and coefficient of variation (CV) $= \sigma_R$ in the simulations.

The cell divides into two symmetrical cells on average (mean $= 1/2$). The inaccuracy in setting the division plane at the end of $n^{th}$ generation of a lineage is $\delta_n$ which has a mean $= 0$ and standard deviation $= \sigma_m$. The length at birth in the $n+1^{th}$ generation ($L_{b,n+1}$) is related to the length at division in the previous ($n^{th}$) generation ($L_{d,n}$) as,

$$L_{b,n+1} = \frac{L_{d,n}}{2}(1+\delta_n) + \frac{R}{3}. \tag{S3}$$

The $\frac{R}{3}$ term in Eq. S3 is added to denote the addition of a hemispherical pole upon the contraction of the division plane while conserving the cell volume.

Throughout our calculations, we will assume that fluctuations in cell width ($\zeta_R(0, \sigma_R)$) and the division noise ($\zeta_s(0, \sigma_{bd})$) are small. On substituting $\alpha = 1$ in Eq. S1- corresponding to volume sizer- and rearranging Eq. S2 by keeping noise terms up to first order, we find the cell length at division to be,

$$L_{d,n} \approx \frac{2V_0}{\pi R_0^2} + \frac{2}{3}R_0 + \frac{2V_0}{\pi R_0^2}\zeta_{s,n} + \zeta_R\left(\frac{2}{3}R_0 - \frac{4V_0}{\pi R_0^2}\right). \tag{S4}$$

Note that $\zeta_{s,n}$ is the value of $\zeta_s$ in the $n^{th}$ generation. The variance of length at division for a population of cells from different lineages is,

$$\sigma_d^2 = \frac{9V^2\sigma_{bd}^2 + 4(\pi R_0^3 - 3V)^2\sigma_R^2}{(3\pi R_0^2)^2}, \tag{S5}$$

where $V = 2V_0$. We assumed that the division noise in a cell cycle is independent of the width fluctuations. Eq. S5 is identical to that in [S6] except for the additional $\sigma_e^2$ corresponding to the variance contribution from measurement errors of cell lengths.

To calculate the birth length, we simplify Eq. S3 keeping the first-order noise terms,

$$L_{b,n+1} \approx \frac{V}{2\pi R_0^2} + \frac{2}{3}R_0 + \frac{V}{2\pi R_0^2}\zeta_{s,n} + \zeta_R\left(\frac{2}{3}R_0 - \frac{V}{\pi R_0^2}\right) + \delta_n\left(\frac{V}{2\pi R_0^2} + \frac{R_0}{3}\right). \tag{S6}$$

Assuming that $\delta_n$ is independent of both $\zeta_{s,n}$ and $\zeta_R$, the variance of the birth length for a cell population is,

$$\sigma_b^2 = \left(\frac{V}{2\pi R_0^2}\right)^2\sigma_{bd}^2 + \left(\frac{2}{3}R_0 - \frac{V}{\pi R_0^2}\right)^2\sigma_R^2 + \left(\frac{V}{2\pi R_0^2} + \frac{R_0}{3}\right)^2\sigma_m^2. \tag{S7}$$

##### S1.1.1 $L_d$ vs $L_b$ plot is consistent with adder model predictions

Next, we will show that such an apparent adder correlation in length can arise due to cell width fluctuations in a volume sizer model (Figure 1 in the main text and Ref. [S6]). The slope of the best linear fit for $L_d$ vs $L_b$ plot is one for the adder model.

The slope of $L_d$ vs $L_b$ plot can be calculated as,

$$m_{bd} = \frac{Cov(L_{b,n}, L_{d,n})}{\sigma_b^2}, \tag{S8}$$

where $Cov(L_{b,n}, L_{d,n})$ is the covariance between $L_b$ and $L_d$.

Using the expressions of $L_{b,n}$ and $L_{d,n}$ from Eqs. S6 and S4, respectively,

$$Cov(L_{b,n}, L_{d,n}) = \left(\frac{2}{3}R_0 - \frac{V}{\pi R_0^2}\right)\left(\frac{2}{3}R_0 - \frac{2V}{\pi R_0^2}\right)\sigma_R^2. \tag{S9}$$

On substituting Eq. S9 into Eq. S8, we obtain,

$$m_{bd} = \frac{\left(\frac{2}{3}R_0 - \frac{V}{\pi R_0^2}\right)\left(\frac{2}{3}R_0 - \frac{2V}{\pi R_0^2}\right)\sigma_R^2}{\left(\frac{V}{2\pi R_0^2}\right)^2\sigma_{bd}^2 + \left(\frac{2}{3}R_0 - \frac{V}{\pi R_0^2}\right)^2\sigma_R^2 + \left(\frac{V}{2\pi R_0^2} + \frac{R_0}{3}\right)^2\sigma_m^2}. \tag{S10}$$

If the variance contribution from measurement errors is included in $\sigma_b^2$, the slope in Eq. S10 will be identical to that in Ref. [S6]. Using values from Ref. [S6] - $V = 3.77\ \mu m^3$; $R_0 = 0.55$ $\mu m$; $\sigma_{bd} = 0.065$; $\sigma_R = 0.035$; $\sigma_m = 0.032$- we obtain the slope to be 0.89. This is very close to the prediction for length adder (slope = 1). Note that measurement errors in length only change the variance of $L_b$ i.e., the denominator in Eq. S10. The slope changes only slightly ($= 0.86$) upon including the measurement error to birth lengths (standard deviation $= \sigma_e =$ $0.04\ \mu m$).

##### S1.1.2 Correlation between birth lengths in successive generations do not agree with experiments

Next, we calculate the correlation between the birth length of the $n^{th}$ cell cycle ($L_{b,n}$) and the birth length $k$ generations later ($L_{b,n+k}$). An ideal cell cycle model should be able to explain the correlations between all cell cycle variables including between $L_{b,n}$ and $L_{b,n+k}$. In this section, we will show that while the $L_d$ vs $L_b$ plot points to a length adder, the correlation between $L_{b,n}$ and $L_{b,n+k}$ does not agree with it when the underlying control is volume sizer.

Similar to Eq. S6, the length at birth $k$ generations after is,

$$L_{b,n+k} \approx \frac{V}{2\pi R_0^2} + \frac{2}{3}R_0 + \frac{V}{2\pi R_0^2}\zeta_{s,n+k-1} + \zeta_R\left(\frac{2}{3}R_0 - \frac{V}{\pi R_0^2}\right) + \delta_{n+k-1}\left(\frac{V}{2\pi R_0^2} + \frac{R_0}{3}\right). \tag{S11}$$

We will quantify correlations between $X$ and $Y$ using Pearson correlation coefficients,

$$r_{xy} = \frac{Cov(x, y)}{\sigma_x \sigma_y}, \tag{S12}$$

where $\sigma_x$ and $\sigma_y$ are the standard deviations of $X$ and $Y$, respectively. At steady state, the standard deviation of $L_{b,n}$ and $L_{b,n+k}$ are equal to $\sigma_b$.

Using Eqs. S6 and S11, we obtain,

$$Cov(L_{b,n}, L_{b,n+k}) = \left(\frac{2}{3}R_0 - \frac{V}{\pi R_0^2}\right)^2 \sigma_R^2. \tag{S13}$$

The Pearson correlation coefficient between $L_{b,n}$ and $L_{b,n+k}$ is,

$$r = \frac{\left(\frac{2}{3}R_0 - \frac{V}{\pi R_0^2}\right)^2 \sigma_R^2}{\left(\frac{V}{2\pi R_0^2}\right)^2 \sigma_{bd}^2 + \left(\frac{2}{3}R_0 - \frac{V}{\pi R_0^2}\right)^2 \sigma_R^2 + \left(\frac{V}{2\pi R_0^2} + \frac{R_0}{3}\right)^2 \sigma_m^2}. \tag{S14}$$

The equation is the same for any value of $k$ for which the radius can be assumed to be constant. Using the values from Ref. [S6], $r = 0.43$ (for $\sigma_e = 0$) and $r = 0.41$ (for $\sigma_e = 0.04$ $\mu m$). We also verify the results using simulations in Figure 2F of the main text.

#### S1.2 Model 2: Length growth at mid-plane before cell division in *Escherichia coli*

The model in Fachetti *et al.* was inspired by cell growth studies in fission yeast [S6]. However, unlike fission yeast, *E. coli* cells do not undergo an abrupt increase in length before birth. *E. coli* cells start constricting at the mid-cell and form the new hemispherical poles before the division event [S10]. In this section, we will explore a cell cycle model where there are two fully formed daughter cells at division. Further, we will relax the assumption that the cell width is the same in a lineage. We will allow the radius to vary during the cell cycle but the radius across generations might still be correlated. The schematic of the cell cycle model motivated here is shown in Figure 3A of the main text.

The division length and birth length are calculated differently as compared to the previous section. According to this model, the cell forms two daughter cells with two hemispherical poles. Thus, the length at division is,

$$L_{d,n} = \frac{V_{d,n}}{\pi R_n^2} + \frac{4}{3} R_n, \tag{S15}$$

where $R_n$ is the cell radius at the time of division of generation $n$.

The positioning of the septum is partly determined by the Min system oscillations which divide the cells, on average, symmetrically in length [S11, S12]. For this discussion, we consider cells dividing into two symmetrical cells, on average, by volume. However, the choice of division by length or volume is irrelevant. We will obtain the same qualitative results if cells were dividing by length as shown in Figure S3. Thus, the volume at birth in the next generation $n + 1$ is,

$$V_{b,n+1} = V_{d,n} \left( \frac{1}{2} + \delta_n \right), \tag{S16}$$

where $\delta_n$ is the noise in volume division ratio $= \frac{V_{b,n+1}}{V_{d,n}}$ for a cell dividing in generation $n$. It has a mean $= 0$ and standard deviation $= \sigma_m$. The length at birth in generation $n + 1$ ($L_{b,n+1}$) is,

$$L_{b,n+1} = \frac{V_{b,n+1}}{\pi R_n^2} + \frac{2}{3} R_n. \tag{S17}$$

The underlying assumption in Eq. S16 is that the radius doesn't change during the division event but can undergo variation during a cell cycle such that the radius at the end of $n + 1^{th}$ generation is,

$$R_{n+1} = R_0 + c(R_n - R_0) + \sqrt{1 - c^2} R_0 \zeta_{R,n+1}. \tag{S18}$$

Here, $R_0$ is the mean radius, $c$ is the correlation between radii in consecutive generations, and $\zeta_{R,n+1}$ is the noise in cell radius. $\zeta_{R,n}$ is assumed to be normally distributed with mean 0, variance $(\langle\zeta_{R,n}^2\rangle) = \sigma_R^2$ and the noise is independent of that in a different generation i.e., $\langle\zeta_{R,n}\zeta_{R,n+1}\rangle = 0$.

On Taylor expanding and keeping the first-order terms, $L_{b,n}$ in Eq. S17 is found to be,

$$L_{b,n} \approx \frac{V_{b,n}}{\pi R_0^2} + \frac{2}{3}R_0 + \left(\frac{2}{3}R_0 - \frac{2V_0}{\pi R_0^2}\right)\zeta'_{R,n-1}, \tag{S19}$$

where $\zeta'_{R,n} = c\frac{R_{n-1}-R_0}{R_0} + \sqrt{1-c^2}\zeta_{R,n}$. Using Eq. S18, we find the covariance, $Cov(\zeta'_{R,n-1}, \zeta'_{R,n}) = c\sigma_R^2$.

Similarly, keeping till the first order terms, $L_{d,n}$ is calculated by substituting Eq. S1 into Eq. S15,

$$L_{d,n} \approx \frac{2(1-\alpha)V_{b,n}}{\pi R_0^2} + \left(\frac{4}{3}R_0 + \frac{2\alpha V_0}{\pi R_0^2}\right) + \frac{2\alpha V_0}{\pi R_0^2}\zeta_{s,n} + \left(\frac{4}{3}R_0 - \frac{4V_0}{\pi R_0^2}\right)\zeta'_{R,n}. \tag{S20}$$

Next, we calculate the slope of the best linear fit of $L_d$ vs $L_b$ plot using Eq. S8. The covariance can be calculated using Eqs. S19 and S20.

$$Cov(L_{b,n}, L_{d,n}) = \frac{2(1-\alpha)\sigma_{vb}^2}{(\pi R_0^2)^2} + 8c\left(\frac{R_0}{3} - \frac{V_0}{\pi R_0^2}\right)^2 \sigma_R^2. \tag{S21}$$

Variance in length at birth which is the denominator in Eq. S8 is,

$$\sigma_b^2 = \frac{\sigma_{vb}^2}{(\pi R_0^2)^2} + 4\left(\frac{R_0}{3} - \frac{V_0}{\pi R_0^2}\right)^2 \sigma_R^2. \tag{S22}$$

The variance in volume at birth $(\sigma_{vb})$ is calculated using Eqs. S1 and S16,

$$\sigma_{vb}^2 = \frac{\alpha^2\sigma_{bd}^2 + 4\sigma_m^2}{\alpha(2-\alpha)}V_0^2. \tag{S23}$$

Hence, we find the slope to be,

$$m_{bd} = \frac{2(1-\alpha)\frac{\alpha^2\sigma_{bd}^2+4\sigma_m^2}{\alpha(2-\alpha)}\frac{V_0^2}{(\pi R_0^2)^2} + 8c\left(\frac{R_0}{3} - \frac{V_0}{\pi R_0^2}\right)^2 \sigma_R^2}{\frac{\alpha^2\sigma_{bd}^2+4\sigma_m^2}{\alpha(2-\alpha)}\frac{V_0^2}{(\pi R_0^2)^2} + 4\left(\frac{R_0}{3} - \frac{V_0}{\pi R_0^2}\right)^2 \sigma_R^2}. \tag{S24}$$

At $n+k^{th}$ generation, the length at birth follows from Eq. S19,

$$L_{b,n+k} \approx \frac{V_{b,n+k}}{\pi R_0^2} + \frac{2}{3}R_0 + \left(\frac{2}{3}R_0 - \frac{2V_0}{\pi R_0^2}\right)\zeta'_{R,n+k-1}. \tag{S25}$$

We want to find the Pearson correlation coefficient between length at birth in $n^{th}$ and $(n+k)^{th}$ generation. The covariance between the two variables is,

$$Cov(L_{b,n}, L_{b,n+k}) = \frac{Cov(V_{b,n}, V_{b,n+k})}{(\pi R_0^2)^2} + 4\left(\frac{R_0}{3} - \frac{V_0}{\pi R_0^2}\right)^2 Cov(\zeta'_{R,n-1}, \zeta'_{R,n+k-1}). \tag{S26}$$

The two covariance terms on the right side of Eq. S26 can be calculated using Eq. S1 and Eq. S18. Substituting the covariance expressions into Eq. S26 and calculating the Pearson correlation coefficient using Eq. S12,

$$r = \frac{(1-\alpha)^k\frac{\alpha^2\sigma_{bd}^2+4\sigma_m^2}{\alpha(2-\alpha)}\frac{V_0^2}{(\pi R_0^2)^2} + 4c^k\left(\frac{R_0}{3} - \frac{V_0}{\pi R_0^2}\right)^2 \sigma_R^2}{\frac{\alpha^2\sigma_{bd}^2+4\sigma_m^2}{\alpha(2-\alpha)}\frac{V_0^2}{(\pi R_0^2)^2} + 4\left(\frac{R_0}{3} - \frac{V_0}{\pi R_0^2}\right)^2 \sigma_R^2}. \tag{S27}$$

The value of $r$ decreases for increasing $k$ as verified using the adder model simulations (Figure S4A). For $\sigma_R << \frac{\alpha^2\sigma_{bd}^2+4\sigma_m^2}{\alpha(2-\alpha)}$, we find that $r(k) \approx (1-\alpha)^k$. Hence, in this regime, we regain the correlation formula for the case where radius variability vanishes. This is evident in Figure S4B where the correlation approaches $(1-\alpha) = 0.5$ for the adder model ($\alpha = 0.5$) when $\sigma_R$ is small compared to $\frac{\alpha^2\sigma_{bd}^2+4\sigma_m^2}{\alpha(2-\alpha)}$.

##### S1.2.1 Cell width for a particular lineage is fixed.

Similar to the model in Section S1.1, we can test a volume sizer ($\alpha$=1) model where the cell width is fixed in a particular lineage ($c = 1$). Substituting $\alpha = c = 1$ in Eq. S27, we get,

$$r = \frac{4\left(\frac{R_0}{3} - \frac{V_0}{\pi R_0^2}\right)^2 \sigma_R^2}{\frac{\alpha^2\sigma_{bd}^2 + 4\sigma_m^2}{\alpha(2-\alpha)}\frac{V_0^2}{(\pi R_0^2)^2} + 4\left(\frac{R_0}{3} - \frac{V_0}{\pi R_0^2}\right)^2 \sigma_R^2}. \tag{S28}$$

Although the exact expression differs from the previous section, we find that the correlation between birth lengths $k$ generations apart is independent of $k$, similar to the previous section and contrary to experimental results.

##### S1.2.2 Using conditional correlations to estimate $\alpha$

We aim to find the underlying volume regulation strategy despite the cell width fluctuations. In the main text, we explain using a graphical representation (Figure 3B), the reason for using the conditional correlation $(L_{b,n}, L_{d,n})|(R_{n-1}, R_n)$ for that purpose. In this section, we explain the method based on the model motivated in the previous section. Then, we estimate the accuracy of the method to elucidate the underlying division strategy.

Eqs. S19 and S20 provide an expression for the birth and division lengths, respectively. Rearranging the radius fluctuation terms, we obtain,

$$L_{b,n} - \frac{2V_0}{\pi R_0^2} - \left(\frac{2}{3} - \frac{2V_0}{\pi R_0^3}\right) R_{n-1} \approx \frac{V_{b,n}}{\pi R_0^2}, \tag{S29}$$

$$L_{d,n} - \left((2+\alpha)\frac{2V_0}{\pi R_0^2}\right) - \left(\frac{4}{3}R_0 - \frac{4V_0}{\pi R_0^2}\right) R_n \approx \frac{2(1-\alpha)V_{b,n}}{\pi R_0^2} + \frac{2\alpha V_0}{\pi R_0^2}\zeta_{s,n}. \tag{S30}$$

The term on the left side of Eq. S29 corresponds to the residual $(L_{b,n}|(R_{n-1}, R_n))$ obtained on linear regression of $L_{b,n}$ on $R_{n-1}$ and $R_n$. Similarly, the left side of Eq. S30 corresponds to the residual $(L_{d,n}|(R_{n-1}, R_n))$ obtained by regressing $L_{d,n}$ instead of $L_{b,n}$. The slope of the best linear fit of $L_{d,n}|(R_{n-1}, R_n)$ vs $L_{b,n}|(R_{n-1}, R_n)$ plot is $2(1-\alpha)$. Thus, this method can be used to obtain the division strategy parameter, $\alpha$.

The expressions in Eqs. S29 and S30 assume that there is no contribution to the cell width fluctuations from measurement noise. However, the measured cell radius ($R_{t,n}$) in the experiments is a combination of the actual radius ($R_n$) and measurement noise in radius ($\zeta_{Re,n}$). Assuming an additive measurement noise in cell radius, we obtain the measured cell radius to be,

$$R_{t,n} = R_0 + c(R_{n-1} - R_0) + \sqrt{1-c^2} R_0 \zeta_{R,n} + \zeta_{Re,n}. \tag{S31}$$

We assume that $\zeta_{Re,n}$ is normally distributed with zero mean and standard deviation $= \sigma_{Re} R_0$. It is also assumed to be independent of $R_n$ but it can be correlated with $\zeta_{Re,n-1}$, i.e., $\langle \zeta_{Re,n-1} \zeta_{Re,n} \rangle = c_e$. Using Eq. S31, we obtain the CV in measured cell radius to be ($\sigma_{Rt}$),

$$\sigma_{Rt}^2 = \sigma_R^2 + \sigma_{Re}^2. \tag{S32}$$

The correlation between radius in consecutive generations is,

$$c_t = \frac{c\sigma_R^2 + c_e \sigma_{Re}^2}{\sigma_R^2 + \sigma_{Re}^2}. \tag{S33}$$

We wanted to test whether the value of $\alpha$ calculated using the conditional correlation $((L_{b,n}, L_{d,n})|(R_{n-1}, R_n))$ is accurate for different values of $\sigma_{Re}$ and $c_e$. We vary the value of $\sigma_{Re}$ and $c_e$ between 0 to $\sigma_{Rt}$ and 0 to 1, respectively, in the simulations of the model shown in Figure 3A of the main text. Further, we restricted ourselves to those values of $\sigma_{Re}$ and $c_e$ for which correlation $c$ was between 0 and 1. We measure the cell volumes, and lengths at birth and division, and the cell radius. We plot the difference between the $\alpha$ values calculated using the correlation between birth and division volumes, and the conditional correlation $((L_{b,n}, L_{d,n})|(R_{n-1}, R_n))$. We find that the difference between the two alpha values is small

- within 0.1 of each other (Figure S5A).

Next, we wanted to test that the conditional correlation $((L_{b,n}, L_{d,n})|(R_{n-1}, R_n))$ is a suitable method to probe cell cycle regulation mechanisms regardless of the underlying cell cycle model. Hence, we tested the conditional correlation method on cell regulation strategies other than the model where the cell division volume is solely determined by the birth volume. In these cell cycle models, the size at the initiation of DNA replication also determines the division size [S1–S3, S13]. We tested two previously proposed models (the CH model and the parallel adder model) where DNA replication solely controls cell division. In one model (CH model), cell division happens after a certain average time from the initiation of DNA replication [S2, S14]. In the parallel adder model, division happens after a certain volume per origin is added from the initiation volume [S13, S15]. Another strategy called the concurrent processes model was also tested where both DNA replication and cell birth control the cell division volume. In this model, cell division happens when the slowest of the two processes - 1. a certain time from initiation of DNA replication elapses, 2. after a particular volume from birth is added- is completed [S1, S3]. In the simulation of these models, we track the birth and division volumes and lengths, and the cell width for multiple generations and lineages. To test the accuracy of the conditional correlation $(L_{b,n}, L_{d,n})|(R_{n-1}, R_n)$ as an indicator of the division volume control strategy in these models, we find the difference between two correlation values - 1. $V_{b,n}$ and $V_{d,n}$, 2. conditional correlation between $L_{b,n}$ and $L_{d,n}$ upon fixing $R_{n-1}$ and $R_n$. The difference between the two correlations for different values of $\sigma_{Re}$ and $c_e$ is small - again within 0.1 of each other (SI Figs. 5B-5D).

Thus, the conditional correlation $(L_{b,n}, L_{d,n})|(R_{n-1}, R_n)$ is an appropriate method to find the underlying cell cycle strategy controlling division volume using cell length data and despite cell width fluctuations.

### S2 Data analysis

#### S2.1 Analyzing experimental data

We analyzed mother-machine data from Refs. [S5], [S4], and [S3]. These datasets were chosen as they had single-cell width measurements along with cell lengths at birth and division. We restricted our analysis to those cells for which we could find the daughter and granddaughter cells. Additionally, only for the experimental datasets in Ref. [S5], we removed the cells whose radii were outliers. These cells appeared as smaller separate islands in the $R_n$ vs $R_{n-1}$ plot and could be identified visually. We used these cells for the calculation of model parameters for six different growth conditions of Ref. [S5] (Table S3) and to find the correlation and conditional correlation involving birth lengths, division lengths, and cell radius (Table S1, Table S2, Table 1 and Figure 4 of the main text).

#### S2.2 Estimating model parameters

In the section "Implications on cell width fluctuations" of the main text, we substitute model parameters into Eq. 3 of the main text to calculate the correlation between birth lengths in successive generations (Table S3). In this section, we explain the methodology to determine various model parameters.

The model parameters were estimated using experimental data from Ref. [S5]. The mean radius ($R_0$), and CV ($\sigma_R$) are the average and CV of the measured radius in a particular experimental dataset. $c$ is the Pearson correlation coefficient of the cell radius between mother and daughter cells. The average volume at birth ($V_0$) is $\pi R_0^2 \langle L_b \rangle - \frac{2}{3}\pi R_0^3$, where $\langle L_b \rangle$ is the mean length at birth. $\sigma_m$ is the standard deviation of the division ratio which is the ratio of the length at birth in the daughter cell and length at division in the mother cell. $\alpha$ is calculated using the slope of the best linear fit of $L_{d,n}|(R_{n-1}, R_n)$ vs $L_{b,n}|(R_{n-1}, R_n)$ plot. Using the estimates of these parameters and Eq. S22, we can calculate the standard deviation of division size noise, $\sigma_{bd}$. Note that in one of the seven conditions studied in Ref. [S5], we obtain a complex value for $\sigma_{bd}$. We do not show that dataset in Table S3.

### S3 Simulations

Simulations were carried out using MATLAB R2021b.

In Figure 2 of the main text, and Figure S1, we carried out simulations of the "Pole formation after division" model. The parameters are the same as in Ref. [S6]. The cell division strategy being used was a volume sizer with the critical volume for division being $2V_0 = 3.77\mu m^3$. 10000 cell lineages were initiated with a particular division volume drawn from a normal distribution with mean size $2V_0$ and CV = 0.065. Each cell lineage was simulated for 25 generations. The cell radius $R$ for each lineage was drawn from a normal distribution with mean = $0.55\mu m$ and CV = 0.035 and it stayed constant over the 25 generations. The results do not change on simulating over 50 generations. The length at division is calculated based on a spherocylindrical cell geometry in Figure 2A of the main text. Upon division, the length at birth, division, and radius were noted for the cell. A measurement error, drawn from a normal distribution with mean 0 and standard deviation = 0.04 $\mu m$, was added to each length measurement. Upon division, we tracked one of the two daughter cells in the next generation. The cells divided, on average, symmetrically by length with a CV = 0.032. Due to the hemispherical pole formation at mid-cell after birth, $\frac{R}{3}$ was added to the length at birth to keep the total volume conserved.

In Figure S2 of the main text, we simulate the "Pole formation before division" model. We have the same initial conditions and parameters as the "Pole formation after division" model. The difference is in the calculation of birth and division lengths. We assume that there are two fully formed cells just before division (Figure S2A). Also, the cell divides, on average, symmetrically by volume with CV = 0.032. We also test the same model with the cell dividing symmetrically by length on average in Figure S3.

In Fig. 3 of the main text, and Figure S4, we simulate the model "Changing radii in a lineage" (Figure 3A). The model parameters are determined using the experimental data in the fastest growth condition ($T_d$ = 17 min) of Ref. [S5]. 10000 cells are initialized with a division volume drawn from a normal distribution with mean, $2V_0 = 5.54 \mu m^3$ and CV = 0.19. The cells have a radius drawn from a normal distribution with mean = 0.49 $\mu m$ and standard deviation = 0.0765. Each lineage is simulated over 25 generations. The radii are correlated (Pearson correlation coefficient = 0.86) between consecutive generations and have the same mean (= 0.49 $\mu m$) and CV (= 0.0765) as the initial cells. In Figure S4B, we vary the CV of cell radius to be between 0 and 0.13. The division and birth lengths are determined as stated in Figure 3A of the main text. The cell divides, on average, symmetrically by volume (CV = 0.06). The division volume is set by an adder model from cell birth with mean volume added = 2.77 $\mu m^3$. The size additive division volume noise has a normal distribution with mean = 0 and standard deviation = 0.53 $\mu m^3$. The birth and division lengths have an additive measurement error which is drawn from a normal distribution with mean = 0 and standard deviation = 0.04 $\mu m$.

In Figure S5, we wanted to verify that the conditional correlation method put forth in Section S1.2.2 is an appropriate method to study the cell cycle regulation mechanisms even when the cell radius is not accurately determined. Thus, we fix the CV of the measured cell radii = (0.0765) and the correlation between the radii in consecutive generations (= 0.86). We vary the contribution of the measurement error to the radius CV and the correlation between the measurement errors in radii measurements of consecutive generations. The actual radius CV and the correlation between the actual radii in consecutive generations are obtained using Eqs. S32 and S33, respectively. Each plot in Figure S5 is obtained using simulations of a different cell cycle model. In Figure S5A, we follow the simulation procedure as mentioned in the previous paragraph ("Changing radii in a lineage" model). In Figure

S5B, we plot the simulation results of the concurrent processes model. The division event is controlled by the slowest of the two processes - 1. The addition of a particular volume (= 1.10 $\mu m^3$) from birth. A size additive division noise, drawn from a normal distribution, is added to the adder model from birth (mean = 0; standard deviation = 0.2 $\mu m^3$), 2. The division happens after a time = 55 min from the initiation of DNA replication. We add a time additive division noise drawn from a normal distribution with mean = 0 and standard deviation = 12 min. In Figure S5C, we plot the results from the CH model simulations. In this model, division happens after a time $T$ from the initiation of DNA replication. $T$ is drawn from a normal distribution with mean = 60 min and CV = 0.05. For Figure S5D, we simulate the parallel adder model where the division event occurs after the addition of $\Delta_{id}$ volume per origin from the initiation of DNA replication. $\Delta_{id}$ is drawn from a normal distribution with mean = 0.5 $\mu m^3$ and CV = 0.1. For all of the models, the initiation of DNA replication happened upon the addition of $\Delta_{ii}$ volume per origin from the initiation in the previous cell cycle. $\Delta_{ii}$ is drawn from a normal distribution with mean = 0.69 $\mu m^3$, CV = 0.1 for the concurrent processes model; mean = 0.7, CV = 0.15 for the CH model and mean = 0.65, CV = 0.2 for the parallel adder model. In the simulations, 10000 cells were initiated at birth with a volume = 0.6545 $\mu m^3$. Each cell grew exponentially with a different growth rate drawn from a normal distribution with mean = 0.0107 $min^{-1}$ and CV = 0.1. Upon division, lengths at birth and division, and the cell radius were noted with measurement errors. For the length measurements, the error was additive and it was drawn from a normal distribution with mean = 0 and standard deviation = 0.04 $\mu m$. We tracked one of the two daughter cells over 25 generations for each lineage. The cells divided, on average, symmetrically in volume with a standard deviation in division ratio = 0.03. The same parameters are also used for the simulations in Figure S6.

### S4 Cell controls surface area instead of volume

Till now we have discussed scenarios where the cell cycle control is on cell volume. However, if we assume that biomass is the relevant quantity being controlled, we need to find a cell geometry characteristic that is most related to biomass accumulation. Ref. [S16] observed that biomass growth is proportional to cell surface area growth during a cell cycle, hence, cell surface area could be a suitable proxy for it. In this section, we discuss a case where the cell surface area is being regulated. We find that cell width fluctuations have a similar effect on the correlations between the lengths at birth and division, and birth lengths $k$ generations apart.

We assume that the cell divides when it reaches a surface area $S_d$ which is solely determined by its surface area at birth,

$$S_d = 2(1-\alpha)S_b + 2\alpha S_0(1+\zeta_s(0,\sigma_{bd})). \tag{S34}$$

The equation is similar to Eq. S1 with cell volume replaced with cell surface area.

Assuming that the cell is a spherocylinder, the total surface area of the cell is always related to its length as,

$$S = 2\pi RL. \tag{S35}$$

The above equation holds even for a cell undergoing constriction at the mid-cell. The length at division in generation $n$ and the birth length in generation $n+1$ is as follows,

$$L_{d,n} = \frac{S_{d,n}}{2\pi R_n}, \tag{S36}$$

$$L_{b,n+1} = \frac{S_{b,n+1}}{2\pi R_n}. \tag{S37}$$

We follow the same assumptions as Section S1.2 to model the cell width fluctuations. We assume that cells do not change their radius when they divide. However, the radius changes during the cell cycle such that the radius when the cell divides in generation $n+1$ is related to the radius in generation $n$ as stated in Eq. S18. The cell divides symmetrically by surface area on average with the standard deviation in division ratio being $\sigma_m$ as before. Since cell length is proportional to the surface area, the length division ratio has the same mean and standard deviation.

Assuming the noise in radius ($\zeta_{R,n}$), division ratio ($\delta_n$), and division size noise ($\zeta_{s,n}$) to be small, we obtain, till first order, the expressions for the birth and division lengths in generation $n$,

$$L_{b,n} \approx \frac{S_{b,n}}{2\pi R_0} - \frac{S_0}{2\pi R_0}\zeta'_{R,n-1}, \tag{S38}$$

$$L_{d,n} \approx \frac{2(1-\alpha)S_{b,n}}{2\pi R_0} + \frac{2\alpha S_0}{2\pi R_0} + \frac{2\alpha S_0}{2\pi R_0}\zeta_{s,n} - \frac{2S_0}{2\pi R_0}\zeta'_{R,n}, \tag{S39}$$

where $\zeta'_{R,n} = c\frac{R_{n-1}-R_0}{R_0} + \sqrt{1-c^2}\zeta_{R,n}$, as previously defined. Using Eqs. S38 and S39, we obtain the slope of the best linear fit of $L_d$ vs $L_b$ plot to be,

$$m_{bd} = \frac{2(1-\alpha)\frac{\alpha^2\sigma_{bd}^2+4\sigma_m^2}{\alpha(2-\alpha)} + 2c\sigma_R^2}{\frac{\alpha^2\sigma_{bd}^2+4\sigma_m^2}{\alpha(2-\alpha)} + \sigma_R^2}. \tag{S40}$$

The correlation between birth lengths $k$ generations apart is,

$$r(k) = \frac{(1-\alpha)^k\frac{\alpha^2\sigma_{bd}^2+4\sigma_m^2}{\alpha(2-\alpha)} + c^k\sigma_R^2}{\frac{\alpha^2\sigma_{bd}^2+4\sigma_m^2}{\alpha(2-\alpha)} + \sigma_R^2}. \tag{S41}$$

Eq. S41 is similar to Eq. 3 of the main text assuming that $\frac{\pi R_0^3}{3} << V_0$. There are two contributions to the length correlations - one from cell surface area regulation strategy, $f(S_b)$, and the other from correlated radii across generations. The same arguments that were presented in the case of cell volume regulation in the main text will apply here for calculating $\alpha$ and estimating the intrinsic cell width variability. Hence, our analysis points to width fluctuations having the same effect on length correlations regardless of the cell characteristics (surface area or volume) under regulation.
